# Balanced cell division is secured by two different regulatory sites in OxyS RNA

**DOI:** 10.1101/2023.10.19.563056

**Authors:** Maya Elgrably-Weiss, Fayyaz Hussain, Jens Georg, Bushra Shraiteh, Shoshy Altuvia

## Abstract

By interfering with cell division, the *Escherichia coli* oxidative stress-induced OxyS small RNA brings about cell cycle arrest thus allowing DNA damage repair. Cell division and cell elongation are opposing functions to the extent that inhibition of cell division requires a parallel inhibition of cell elongation for the cells to survive. In this study, we report that in addition to cell division OxyS inhibits *mepS* encoding an essential peptidoglycan endopeptidase responsible for cell elongation. Furthermore, a phylogenetic evolutionary target conservation analysis of OxyS homologs to identify OxyS most common function revealed that the majority of OxyS targets belong to the category of “cell cycle” followed by “peptidoglycan metabolism”, especially cell elongation. *mepS* is the most frequent target of this category, predicted to be regulated by OxyS in 64 % of the 146 organisms studied. We suggest that cell cycle arrest and balancing between cell division and cell elongation are important and conserved functions of the oxidative stress induced sRNA OxyS.

## INTRODUCTION

The *Escherichia coli* oxidative stress induced OxyS sRNA has been proposed to play a role in protecting cells against the damaging effects of mutagens such as hydrogen peroxide and alkylating agents (Altuvia et al. 1997) while overexpression of OxyS has been found to be toxic. Studying the molecular mechanism mediating the protective function of OxyS and/or the toxic phenotype we showed that the antimutator function of OxyS is actually the other side of the coin of its cytotoxic phenotype (Barshishat et al. 2018). By repressing the expression of the essential transcription termination factor *nusG*, OxyS enables read-through transcription into a cryptic prophage encoding *kilR*. The KilR protein interferes with the function of the major cell division protein FtsZ, thus imposing growth arrest. This transient growth inhibition facilitates DNA damage repair, enabling cellular recovery, thereby increasing viability following stress. Namely, by indirectly inhibiting cell division OxyS buys the cell more time to properly repair its DNA. Once the oxidative stress has been resolved, OxyS is no longer expressed and the cell returns to its normal life cycle (Barshishat et al. 2018).

Here we show that OxyS sRNA represses expression of *mepS* encoding peptidoglycan (PG) endopeptidase in addition to the transcription termination factor *nusG*.

MepS promotes cell elongation by making space for the insertion of new PG material (Singh et al. 2015; Truong et al. 2020). PG is an essential cross-linked macromolecule that confers bacterial cell shape and rigidity. In order to maintain the integrity of the PG during its expansion, cross-link cleavage by hydrolases such as MepS must be tightly coupled with cross-link formation catalyzed by synthases. As cell division and cell elongation are contrasting functions, mutants impaired in cell division are sensitive to increased activity of peptidoglycan endopeptidases (Truong et al. 2020). Thus, it makes perfect sense that inhibition of cell division by OxyS is accompanied by repression of *mepS* expression.

## RESULTS AND DISCUSSION

### Base-pairing between OxyS loop A and *mepS* mRNA inhibits expression of *mepS*

Searching for OxyS targets using CopraRNA (Wright et al. 2013), we noticed *mepS* (<u>m</u>urein <u>e</u>ndope<u>p</u>tidase, formerly Spr). *mepS* encoding an essential PG endopeptidase influences bacterial morphogenesis by incorporating new murein (Singh et al. 2015; Truong et al. 2020). As OxyS repression of *nusG* results in inhibition of cell division (Barshishat et al. 2018) and mutants impaired in cell division are sensitive to increased activity of peptidoglycan endopeptidases (Truong et al. 2020), we wondered whether OxyS controls expression of *mepS* in addition to *nusG*.

Based on the CopraRNA program, 11 nucleotides at the 5’-end of OxyS (part of loop A) were predicted to base-pair with the ribosome binding site of *mepS* mRNA (Fig. 1AB). To examine the effect of OxyS on *mepS* expression, we constructed a *mepS*-*lacZ* translation fusion, P*lac*-*oxyS* wild type and two OxyS mutants; P*lac*-*oxyS* Δ15-33 and P*lac*-*oxyS* C18G C30G in which the binding site to *mepS* was deleted or disrupted, respectively (Fig. 1AB). β-galactosidase assays detected a 4.6-fold decrease in *mepS*-*lacZ* expression in the presence of wild type OxyS, while OxyS mutants (Δ15-33 and C18G C30G) failed to repress *mepS* expression (Fig. 1C).

**FIGURE 1.**
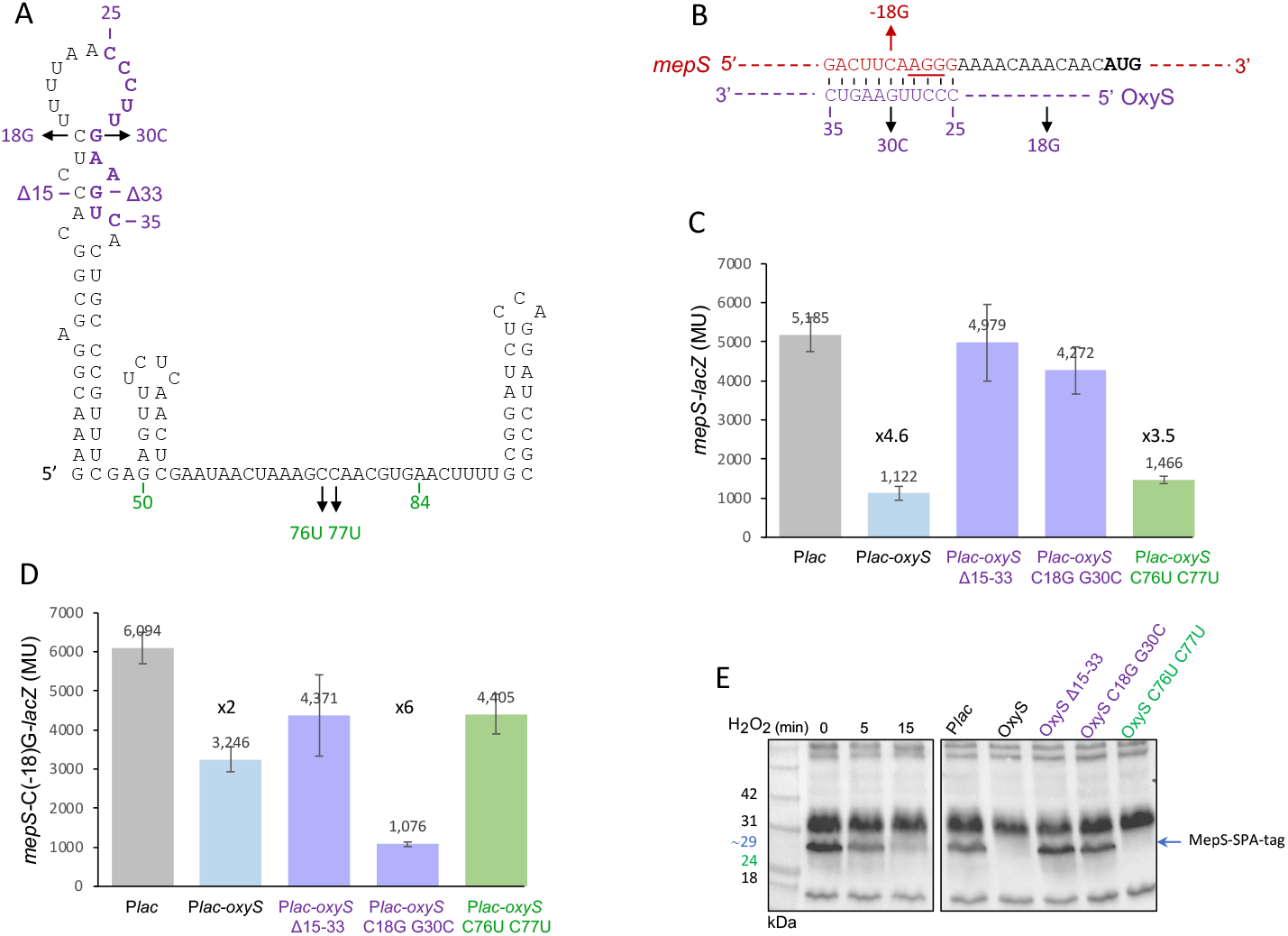
OxyS represses *mepS* expression. (A) Nucleotides 50 to 84 (marked in green) in OxyS are involved in *nusG* binding, while nucleotides 25 to 35 (purple) bind *mepS*. In OxyS Δ15-35 mutant nts 15 to 35 were deleted. In OxyS C18G G30C, the mutation G30C complements (−18)G in *mepS* while C18G restores the disrupted OxyS structure (B) Predicted base pairing between *mepS* (red) and OxyS (purple) RNAs. Indicated are the initiation codon (bold) and the Shine-Dalgarno sequence of *mepS* (underlined). The position of *mepS* and OxyS complementary mutations is indicated. (C, D) Cultures; (MG1655 *mal*::*lacI*^q^Δ*lacZ*::*Tn*10) carrying *mepS-lacZ* wild type and *mepS* C(−18)G*-lacZ* (pSC101*) translational fusion mutant and P*lac*-OxyS wild type and mutants were treated with IPTG (1mM) at OD600 of 0.1-0.2. β-galactosidase activity was measured 60 min after treatment. Results are displayed as mean of 3-6 biological experiments ± standard deviation). (E) To detect the effect of OxyS on MepS protein levels, the sequential peptide affinity (SPA) tag (Zeghouf et al. 2004) was inserted into the MG1655 (*mal*::*lacI*^q^) chromosome adjacent to the carboxy-terminal amino acid of MepS. At OD600 of 0.5 the cultures were exposed to 1 mM H2O2 for the indicated time points whereas cultures carrying OxyS plasmids were treated with 1 mM IPTG for 30 min to induce OxyS expression.

Inhibition of *nusG* by OxyS involves nucleotides 50 to 84 in OxyS. As inhibition of *mepS* by OxyS involves nucleotides 25 to 35, repression of *mepS* by P*lac-oxyS* C76U C77U mutant that is unable to repress *nusG* but carries an intact *mepS* recognition domain was similar to that of wild type OxyS. Repression of *mepS*-*lacZ* carrying C(−18)G, the complementary mutation to OxyS C18G C30G was restored when basepairing was restored (Fig. 1D).

To detect the effect of OxyS on MepS protein levels, the sequential peptide affinity (SPA) tag (Zeghouf et al. 2004) was inserted into the MG1655 (*mal*::*lacI*^q^) chromosome adjacent to the carboxy-terminal amino acid of MepS. At OD600 of 0.5 the cultures were exposed to 1 mM H_2_O_2_ for 5 and 15 min to induce OxyS transcription from the chromosome whereas cultures carrying OxyS plasmids were treated with 1 mM IPTG for 30 min to induce plasmid-encoded OxyS. Western of MepS-SPA showed that upon exposure to H_2_O_2_ MepS-SPA levels decreased with time (Fig. 1E, left panel). Similarly, MepS-SPA levels decreased dramatically in the presence of wild type OxyS and OxyS C76U C77U while OxyS Δ15-33 and OxyS C18G C30G in which the binding site to *mepS* was deleted or disrupted, had no effect on MepS protein levels (Fig. 1E, right panel)

### Short *mepS* mRNA is more responsive to OxyS regulation

Genome-wide studies identified two promoters upstream of *mepS* (Fig. 2A) (Mendoza-Vargas et al. 2009; Chung et al. 2013). Folding of the two mRNA species produced by the two promoters showed that the site to which OxyS binds in the long *mepS* mRNA molecule (G146-G156) is predicted to be sequestered, while the same site in the short molecule (G37-G47) seems more accessible (Figure S1). We examined the effect of OxyS on the short *mepS* transcript using a (ΔP2)P1-*mepS-lacZ* fusion in which the upstream promoter (P2) was deleted leaving the downstream promoter intact and on the long transcript using P2□ΔP1)*-mepS-lacZ* in which the - 10 and -35 sites of downstream promoter (P1) were mutated leaving the upstream promoter intact (Fig. 2A). β-galactosidase assays detected an 8-fold reduction in *mepS*-*lacZ* produced by the short mRNA due to OxyS (Fig. 2B) while repression by OxyS of *mepS*-*lacZ* produced by long transcript resulted in a 3-fold reduction only (Fig. 2C), indicating that *mepS* expression produced by the short transcript is more readily regulated by OxyS. Furthermore, *mepS-lacZ* produced by the long transcript is less accessible to the translation machinery as indicated by the 5.6-fold decrease in β-galactosidase basal activity levels.

**FIGURE 2.**
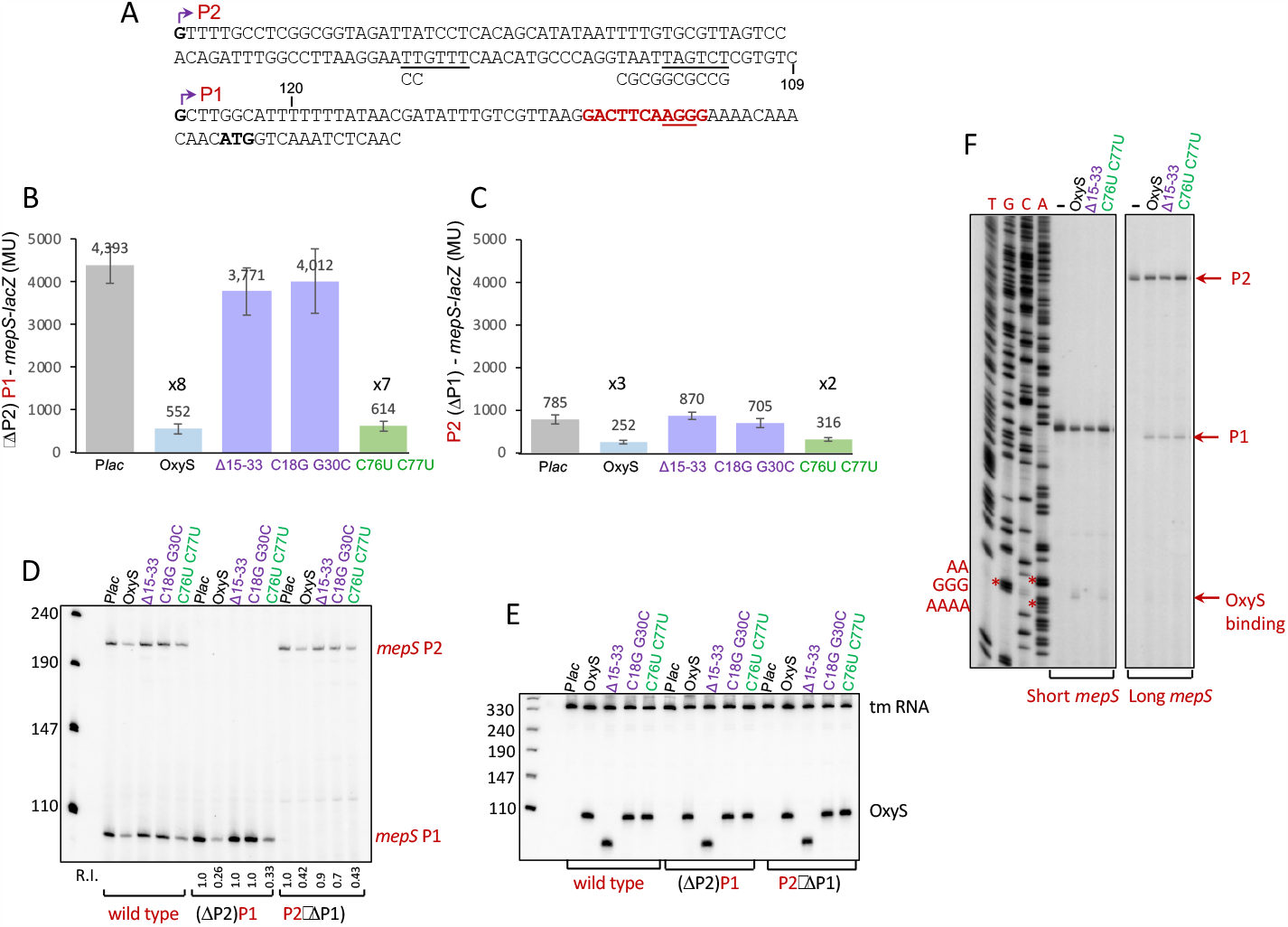
Short *mepS* transcript is responsive to OxyS regulation. (A) *mepS* is transcribed from two promoters P2 and P1 producing long and short mRNAs. To inactivate the downstream promoter P1, the sites -10 and -35 (underlined) were mutated as shown beneath the sequence. The initiation codon (bold) and the Shine-Dalgarno sequence of *mepS* (underlined) are indicated. The sequence in *mepS* that can bind OxyS is in red. (B, C) OxyS affects expression of short *mepS* mRNA (*lacZ* assays). Cultures (MG1655 *mal*::*lacI*^q^□Δ*lacZ*::*Tn*10) carrying (ΔP2)P1*-mepS-lacZ* and P2□ΔP1)*-mepS-lacZ* (pSC101*) translational fusion and P*lac*-OxyS wild type and mutants were treated with IPTG (1mM) at OD600 of 0.1-0.2. β-galactosidase activity was measured 60 min after treatment. Results are displayed as mean of 4-5 biological experiments ± standard deviation). (D) OxyS-*mepS* RNA interaction *in vivo*. Cultures (MG1655 *mal*::*lacI*^q^) carrying OxyS plasmids and chromosomally encoded wild type full-length, short (ΔP2)P1 and long (P2□ΔP1) *mepS* transcripts were treated with IPTG (1mM) at OD600 of 0.2-0.3 for 30 min. The cDNA products generated by primer extension using 30 μg of total RNA and end-labelled *mepS* specific primer 3412 were analyzed in 6% acrylamide 8M urea-sequencing gel alongside with pUC19-MspI labelled marker. Relative intensity (R.I) Band intensities were determined by the ImageLab program. *mepS* mRNA levels were normalized to tm RNA loading control. *mepS*/Plac was used as a 100% reference. (E*) In vivo* RNA samples (10 μg) as in D were separated using 6% urea-polyacrylamide gels (northern blot). The membranes were probed with end-labeled OxyS (3708) and tm RNA (1912) specific primers. tm RNA serves as a loading control. (F) OxyS-*mepS* RNA binding *in vitro. In vitro* synthesized (0.05 pmol) short (139 nt) and long (248 nt) *mepS* mRNAs incubated with and without (5 pmol) synthesized OxyS RNAs (173 nt wild type and 158 nt Δ15-33) at 25°C for 15 min. Primer extension was carried out for 7.5 min at 37°C using end-labeled *mepS* specific primer (3740). The products were analyzed in 6% acrylamide 8M urea-sequencing gel alongside with sequencing reactions.

*In vivo* RNA analysis of chromosomally encoded short ((ΔP2)P1) and long (P2□ΔP1)) *mepS* mRNA constructed by the scarless mutation methodology showed that wild type OxyS expression results in approximately 4 and 2-fold decrease in *mepS* short and long mRNA levels, respectively. Both OxyS mutants with disrupted complementarity to *mepS* have no effect on *mepS* RNA levels, while expression of OxyS C76U C77U that is unable to repress *nusG* but capable of repressing *mepS* reduces *mepS* transcript levels similar to OxyS wild type (Fig. 2D). As the plasmid encoded wild type OxyS levels were similar to the levels of OxyS mutants further indicated that repression of *mepS* is due to *mepS*-OxyS interaction (Fig. 2E).

Given that stable RNA hybrids may block the elongation by reverse transcriptase, we examined the interaction of *in vitro* synthesized OxyS, with short and long *mepS* RNAs using primer extension assays. These assays showed that the interaction of OxyS and OxyS C76U C77U with short *mepS* mRNA results in a termination signal that maps to the site of complementarity (Fig. 2F). No termination signals were detected when OxyS Δ15-33 was incubated with short mRNA or upon incubation of wild type OxyS with the long *mepS* RNA (Fig. 2F).

### Simultaneous inhibition of cell division and cell elongation by OxyS promotes cell recovery

Given that cell division and cell elongation are contrasting functions, mutants impaired in cell division are sensitive to increased activity of peptidoglycan endopeptidases (Truong et al. 2020). Here we show that OxyS represses expression of *mepS* responsible for cell elongation in addition to cell division inhibition. The effect of *mepS* repression on growth was examined using an OxyS mutant in which the complementary sequence to *mepS* was deleted. Cells carrying P*lac-oxyS* Δ15-33 were more inhibited than cells expressing wild type OxyS (Figure 3A) suggesting that decreasing *mepS* expression is advantageous when cell division is inhibited. As shown previously the growth of cells expressing P*lac-oxyS* C76U C77U mutant that is unable to repress *nusG* was similar to that of the control (Barshishat et al. 2018).

**FIGURE 3.**
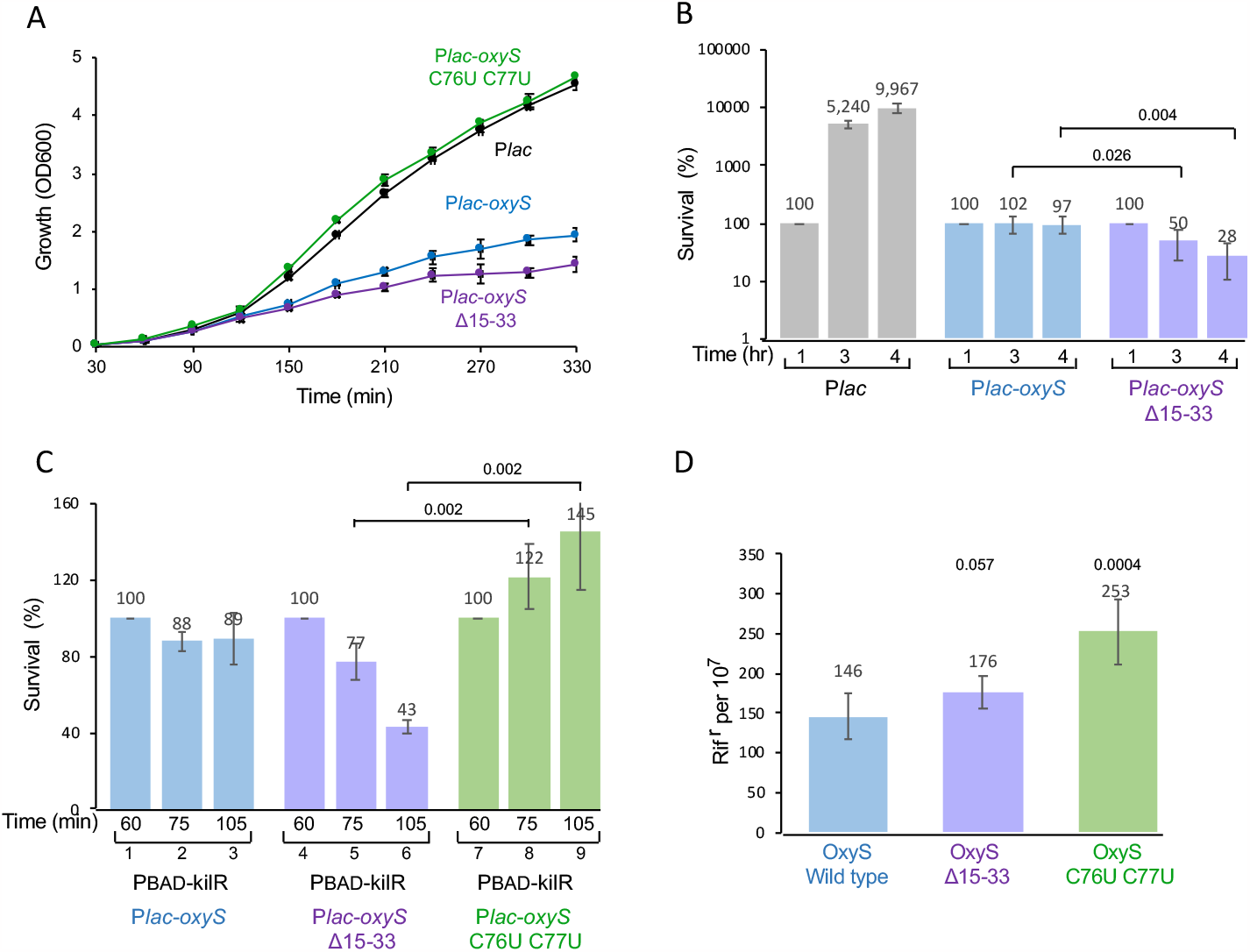
Simultaneous inhibition of cell division and cell elongation by OxyS promotes cell recovery. (A) *E. coli* (MG1655 *mal*::*lacI*^q^) carrying P*lac*-OxyS wild type and mutants were treated with 1 mM IPTG at dilution, growth (OD600) was measured as indicated. (B) Cultures (MG1655 *mal*::*lacI*^q^) carrying control and OxyS plasmids and chromosomally encoded wild type short (ΔP2)P1 *mepS* transcript that was constructed using the scarless mutations methodology were treated with IPTG (1mM) at dilution, CFU were determined 1, 2 and 4 hours after dilution. (C) Cultures (MG1655 *mal*::*lacI*^q^) carrying PBAD-kilR, OxyS plasmids were treated with arabinose 0.2% and IPTG (1mM) at dilution, CFU were determined as indicated in the figure. Results are displayed as mean of 5 biological experiments ± standard deviation). (D) Cultures of wild type OxyS and two chromosomally encoded OxyS mutants (Δ15-33 and C76U C77U) were treated with 0.2 mM of H_2_O_2_ for 30 min to induce OxyS expression and then exposed to 10mM H_2_O_2_ to induce mutations. CFU and rifampicin resistant cells were monitored after 18 hours of growth. Results are displayed as mean of 6 biological experiments ± standard deviation).

Similarly, cell survival assay of strains carrying chromosomally-encoded short *mepS* mRNA and plasmids encoding wild type OxyS and OxyS Δ15-33 revealed that while cells carrying the control plasmid continued to grow and divide, cells expressing OxyS stopped dividing and cells expressing the OxyS Δ15-33 showed a decrease in the number of CFU, indicating that stopping cell division without stopping cell elongation is highly detrimental (Fig. 3B).

By repressing the expression of the essential transcription termination factor *nusG*, OxyS enables read-through transcription into a cryptic prophage encoding *kilR*. The KilR protein interferes with the function of the major cell division protein FtsZ, thus imposing growth arrest (Barshishat et al. 2018). To characterize the effect on cell survival of OxyS regulation of *mepS* separately from *nusG*, we expressed KilR *in trans* from an inducible promoter (PBAD-*kilR*). Plasmid-encoded *in trans* expression of *kilR* accompanied by chromosomally-encoded *kilR* resulted in a dramatic decrease in cell survival in the absence of *mepS* repression (compare lanes 2 and 3 to 5 and 6; OxyS Δ15-33) (Fig. 3C) whereas, *mepS* repression by OxyS mutant lacking the ability to induce chromosomally-encoded *kilR*, rescues cells from the toxic effects of plasmid-encoded *kilR* (compare lanes 2 and 3 to 8 and 9) further confirming the importance of OxyS regulation of both functions together for cells to survive.

Previously, we showed that by inhibiting cell division OxyS buys the cells more time to properly repair its DNA (Barshishat et al. 2018). We tested the relative effect of cell division versus cell elongation on DNA damage repair by monitoring mutation rate in cultures carrying wild type OxyS and two chromosomally encoded OxyS mutants (Δ15-33 and C76U C77U) constructed using the scarless mutations methodology. The cultures were first treated with 0.2 mM of H_2_O_2_ for 30 min to induce OxyS expression and then exposed to 10 mM H_2_O_2_ to induce mutations. Inhibition of cell division only while cell elongation is active (OxyS Δ15-33) results in a slight increase (1.2-fold) in mutation frequency compared to wild type OxyS which inhibits both functions. In contrast, inhibiting cell elongation while cell division is active (OxyS C76U C77U) results in a more pronounced increase (1.7-fold) in the rate of mutations (Fig. 3D).

### Simultaneous inhibition of cell division and cell elongation by OxyS is phylogenetically conserved

To find out what is the most common function of OxyS and to learn whether it involves linking between cell division and cell elongation, we conducted a phylogenetic study of OxyS homologs. A total of 1340 OxyS homologs were detected with the GLASSgo tool in the NCBI nt database (Lott et al. 2018). A phylogenetically weighted sequence logo based on an alignment of all detected homologs shows that OxyS 5’ part is rather variable while the internal region and the Rho-independent terminator are rather conserved (Supplemental Fig. S3). We selected 146 representative homologs (Supplemental Fig. S2) that cover the whole phylogenetic distribution of OxyS for an evolutionary target conservation analysis based on CopraRNA (Wright et al. 2013). For each organism we selected 14 partners for the comparative CopraRNA prediction and performed a gene set enrichment analysis on the top 200 predicted targets (Supplemental Fig. S4 and Table S1). Strikingly, the most frequent category was “cell cycle” that was enriched in 121 of the 146 organisms containing cell cycle and cell division targets including *mraY* (77/146), *murG* (72/146), *nagZ* (70/146), *tolB* (65/146), *zapC* (68/146) and *ftsZ* (54/146). Other categories such as membrane organization, peptidoglycan metabolic process and extracellular polysaccharide biosynthetic process were also frequently enriched, indicating a possible role of OxyS in cell elongation and membrane growth. Widely conserved predicted targets from these categories are e.g *bamA* (124/146), *bamD* (60/146) or *mepS* (97/146).

To test the validity of these predictions, we chose *zapC, ftsZ, murG, minD, bamD* and *mepS* of *Salmonella* for further investigation. Among genes involved in cell division we found *zapC* and *ftsZ lacZ* fusions to be affected by OxyS (Supplemental Fig. S5). Intriguingly, *fts-lacZ* regulation by *Salmonella* OxyS is mediated by the same region that is involved in the regulation of *nusG* by *E. coli* OxyS. Thereby, wild type OxyS of *E. coli* represses expression of *Salmonella ftsZ*-*lacZ* and OxyS mutants that fail to inhibit *nusG* also fail to inhibit *ftsZ-lacZ*. In case of *zapC*, although *zapC-lacZ* is regulated by the 5’ domain of the site involved in *nusG* regulation (position 52 to 66) (Barshishat et al. 2018) *E. coli* OxyS likely fails to regulate *Salmonella zapC* because the sequence of the interaction site differs between *Salmonella* and *E. coli*, which may indicate independent target evolution. The *mepS* target and the interaction site is conserved between *Salmonella* and *E. coli*. Thus, *E. coli* OxyS represses *Salmonella mepS* and OxyS mutants that fail to inhibit *mepS* of *E. coli* also fail to repress *mepS* of *Salmonella* (Supplemental Fig. S5).

### Fluorescence microscopy imaging of cell morphology

We visualized the effect of the separation between cell division and cell elongation on their morphology by fluorescence microscopy. Surprisingly, we found that inhibition of cell division only, while cell elongation is active (OxyS Δ15-35) results in a unique morphology where cells cluster together and adhere to each other (Fig. 4). Given that the morphology of *E. coli* cells expressing KilR which is known to interfere with cell division was similar to that of cells expressing OxyS Δ15-35 mutant indicates that “clumping” results from an imbalance between division and elongation. The morphology of *Salmonella* expressing wild type OxyS appears to be different from the morphology detected in *E. coli*, possibly because the effect of *E. coli* OxyS on cell division is indirect, mediated by KilR that interferes with the function of FtsZ, while in *Salmonella*, OxyS decreases *ftsZ* expression directly conceivably by base paring (Supplemental Fig. S6).

**FIGURE 4.**
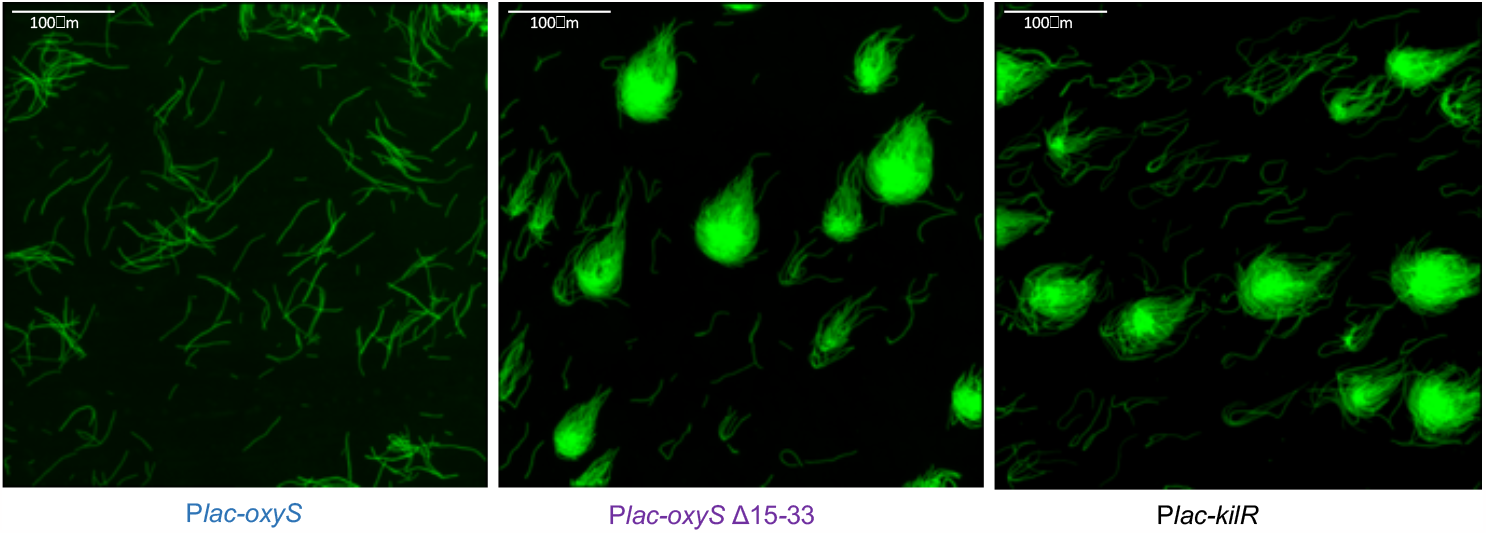
Fluorescence microscopy images of *E. coli* expressing OxyS or *kilR*. Cultures of *E. coli mal*::*lacI*^q^ expressing OxyS wild type and mutant or *kilR* as indicated were grown for 3 hours in the presence of IPTG from dilution. Scale bar 100μm.

In conclusion, in this study we show that *E. coli* OxyS regulates two different target genes associated with cell division using two separate sites: one site interferes with the function of *ftsZ* by inhibiting expression of *nusG*, whereas the second site inhibits expression of *mepS* to prevent cell elongation. By inhibiting both cell division and cell elongation, OxyS rescues cells arrested in division from the destructive consequences of increased activity of peptidoglycan endopeptidases and buys the cell more time to properly repair its DNA.

A phylogenetic evolutionary target conservation analysis of OxyS homologs to identify OxyS most common functions revealed that the majority of OxyS targets belong to the category of “cell cycle” (121/146) followed by “peptidoglycan metabolism” (82/146). Notably, while the involvement of OxyS in cell cycle arrest appears to be broadly conserved, in many of the cases it is achieved by regulating different sets of genes. In *Salmonella*, OxyS affects cell cycle by regulating at least three cell cycle-related genes *ftsZ, zapC* and *mepS* while in *E. coli* cell cycle arrest is achieved through regulation of *nusG* and *mepS*. Intriguingly, *ftsZ* regulation by *Salmonella* OxyS is mediated by roughly the same region that is involved in the regulation of *nusG* by *E. coli* OxyS. The regulation of *nusG* is likely a more recent evolutionary adaption in the genus *Escherichia*. While the *nusG* UTR is largely conserved between all 146 investigated organisms, only *Escherichia* specific changes in the rather un-conserved loop b in OxyS enabled its regulation. In contrast the *mepS*-OxyS interaction site is conserved between *Salmonella* and *E. coli*.

Hundreds of bacterial sRNAs with diverse metabolic functions have been identified so far. However, the number of sRNAs known to influence cell division, either directly or indirectly is still scarce. One example is the *E. coli* prophage encoded DicF sRNA that was discovered more than two decades ago as an inhibitor of cell division (Faubladier et al. 1990). DicF prevents assembly of the septal ring structure by *ftsZ* by repressing translation of *ftsZ* mRNA (Balasubramanian et al. 2016). The sRNA StsR of *Rhodobacter sphaeroides* is induced upon stress conditions and during stationary phase by alternative sigma factors. Expression of StsR provides a regulatory link between cell division and environmental cues (Grützner et al. 2021). EcpR1 is an example of an sRNA that modulates cell cycle. High-level expression of the sRNA EcpR1 in the plant-symbiotic *Sinorhizobium meliloti* has resulted in cell elongation (Robledo et al. 2015).

Exploring the RIL-Seq data for possible association between sRNAs and genes involved in cell division predicts several putative interactions between CyaR sRNA and genes of cell division of which none were confirmed (Melamed et al. 2016). Unlike the above, a search for RprA sRNA partners, led to the discovery that translation of *cpoB* mRNA encoding a cell division coordinator was reduced by 50% in the presence of RprA (Lalaouna et al. 2018).

Most sRNAs control the expression of multiple targets with complementary functions that together compose a mini-regulon. For example, the small RNA RyhB is specifically transcribed under iron-depleted conditions and regulates expression of a large set of target mRNAs, many of which encode iron-containing and iron-storage proteins and are down-regulated by RyhB (Masse and Gottesman 2002; Massé et al. 2005). Likewise, GcvB sRNA controls expression of a large number of functionally related mRNAs whose encoded proteins belong to the class of ABC transporters involved in amino acid metabolism including uptake and biosynthesis (Urbanowski et al. 2000; Sharma et al. 2007; Pulvermacher et al. 2009a; Pulvermacher et al. 2009b; Pulvermacher et al. 2009c; Sharma et al. 2011). In response to glucose-phosphate stress, SgrS sRNA down-regulates the expression of sugar transporters and up-regulates phosphatase *yigL* that dephosphorylates the accumulated sugars to facilitate their export (Vanderpool and Gottesman 2004; Rice and Vanderpool 2011; Sun and Vanderpool 2013). McaS, a mid-stationary phase sRNA regulated by carbon source availability, represses *csgD*, a key player in biofilm formation and activates expression of *flhD* and *pgaA* affecting motility and biofilm respectively. Thus, creating a mini regulon that dis-favors biofilm formation and promotes motility (Jørgensen et al. 2012; Thomason et al. 2012). Slightly different from the above, OxyS controls expression of two linked functions where changes in one function require concomitant changes in the other in order for the cells to survive.

Finally, cell morphology images detected by fluorescence microscopy are striking. Cells in which cell division is inhibited while cell elongation is active display a unique fireball-like morphology. Truong et al showed that filamentous Δ*ftsP* cells overproducing MepS display half the number of normal Z-rings relative to wild-type cells carrying a control plasmid thus interfere with the assembly of mature divisomes (Truong et al. 2020). The phase-contrast images of mutant strains defective for division with *in trans* overproduction of MepS are somewhat reminiscence of the images detected with overproduction of kilR or OxyS Δ15-35. Whether these “fireballs” are formed as a result of an unfavorably high expression of MepS in the context of cell division inhibition and what is the mechanism that leads to their formation remains to be seen.

## MATERIALS AND METHODS

### Bacterial growth conditions

*Escherichia coli* and *Salmonella* cultures were grown at 37°C (200 rpm) in LB medium (pH 6.8). Ampicillin (100 μg/ml), tetracycline (10 μg/ml), chloramphenicol (20 μg/ml), and kanamycin (40 μg/ml) were added where appropriate. P*lacO* promoter was induced with isopropyl β-D-thiogalactoside (IPTG; 1 mM), PBAD promoter was induced with 0.2% arabinose as indicated. Strains, plasmids and primers used in this study are listed in Supplemental Tables S2, S3, and S4.

### Strain construction

To construct Δ*oxySF*::kan, the chromosomal region flanked by genome coordinates 4158287and 4158395 (GenBank entry NC_000913.3) was replaced by the *kan* gene from pKD4 plasmid, using primers 3651 and 3652 as described (Datsenko and Wanner 2000). *oxyS* gene disruption was examined by PCR using flanking primers, 2026 and 2027. To construct *Salmonella* (SL1344 Δ*hisG*::*lacI*^q^:Cm) a PCR fragment carrying P*lacI*-*lacI*^q^ between AatII and XhoI and downstream of cm cassette was amplified using primers 2156 and 2157. The fragment was inserted into *hisG46* mutant of *Salmonella*. Chromosomal scarless point mutations in *mepS* and *oxyS* were carried out as described (Li et al. 2013). Briefly, the *tetA-sacB* cassette from the XTL634 strain chromosome was amplified using the primer pairs 3425-3420 (*oxyS* Δ15-33), 3640-3641 (*oxyS* CC76,77UU), 3722-3723 (*mepS* (ΔP2)) and 3728-3729 (*mepS* (ΔP1)). The PCR products were inserted into MG1655 *mal*::*lacI*^q^ carrying pKD46 plasmid generating insertions of *tetA-sacB* in the selected genes. Next, the products of Gibson reactions carried out on two PCR fragments generated using primer pairs 3427-3428 and 3429-3430 to construct *oxyS* Δ15-33; 3427-3642 and 3643-3430 to construct *oxyS* CC76,77UU; 3726-3724 and 3725-3727 to construct *mepS* (ΔP2) as well as 3730-3726 and 3731-3727 to construct *mepS* (ΔP1) were used to transform the various *tetA-sacB* strains. Colonies sensitive to tetracycline were selected on fusaric acid-containing plates. The newly chromosomally generated scarless mutations were verified by sequencing. MepS-SPA fusion in the chromosome, was constructed using primers (3751 and 3752) designed to amplify the sequential peptide affinity (SPA) tag together with the kanamycin resistance cassette from plasmid pJL148 (Zeghouf et al. 2004). The PCR products were gel purified and then transformed into DY378 cells that were grown at 30°C to OD600 of 0.5 then transferred to 42°C for 15 min. Insertions were confirmed by PCR using primers 3753 and 3691. The products were sequenced using primer 2227. The fusions were transferred into MG1655 *mal*::*lacI*^q^ by P1 transduction.

### Plasmid construction

To construct P*lac*-*oxyS* Δ15-33 and P*lac*-*oxyS* C18G G30C whole plasmid PCR was carried out using P*lac*-*oxyS* plasmid as template and primers 3419-3420 and 3745-3746, respectively. To construct *mepS*-*lacZ* translation fusion pBOG552 (Hershko-Shalev et al. 2016) (Supplemental Table S3), *mepS* fragments were PCR amplified using primers 3411-3412 and chromosomal wild type and *mepS* mutants (ΔP1, ΔP2) as templates. The PCR products were then cloned into the EcoRI and BamHI sites of pBOG552. To construct *mepS*-C(−18)G-*lacZ* fusion the EcoRI and BamHI fragment was inserted into pGEM3 plasmid and subjected to whole plasmid PCR using primers 3754-3755 of which 3754 carries C(−18)G. The mutated fragment was then inserted into pBOG552. To construct PBAD-*kilR, kilR* sequence was amplified from MG1655 chromosomal DNA using primers 3922 and 2701 and cloned into the PstI and HindIII restriction sites of pEF21 plasmid.

To construct *Salmonella mepS* and *zapC*-*lacZ* translation fusions, PCR fragments amplified using primers 3940-3941 and 3942-3943, respectively were cloned into the EcoRI and BamHI sites of pBOG552. To construct P*lacO-ftsZ*-*lacZ* translation fusion the P*lacO*, PCR fragment amplified using primers 3319-3320 and pBR P*lacO* as template was cloned into EcoRI and KpnI. *Salmonella ftsZ* fragment was PCR amplified using primers 3952-3953 and cloned into the KpnI and BamHI sites.

### β-galactosidase assays

Overnight cultures of MG1655 *mal*::*lacI*^q^□Δ*lacZ*::*Tn*10, carrying *mepS-lacZ* translational fusion P*lac* and P*lac*-*oxyS* plasmids (Supplemental Table S3), were diluted 1/100 in 10ml LB medium containing ampicillin and kanamycin and grown to OD600 of 0.1-0.2. Thereafter, the cultures were treated with IPTG (1mM) to induce transcription of OxyS. β-galactosidase activity was measured 60 min after IPTG induction. Cultures of *Salmonella* (SL1344 Δ*hisG*::*lacI*^q^:cm) carrying *lacZ* fusion plasmids were induced by IPTG (1 mM) at dilution. Samples were taken 150 min after dilution.

### Western of SPA tagged *mepS*

Overnight cultures of SPA-tagged strain were diluted 1/100 and grown shaking (200 rpm) at 37°C in LB. H2O2 (1mM) was added at OD600 of 0.5 as indicated in the figure. Expression of OxyS was induced by 1mM IPTG, 1h after dilution for 30 min. Pellets were resuspended in 1X Laemmli sample buffer, heated at 95°C for 5 min. and analyzed on SDS-PAGE (10%). The proteins were transferred to a nitrocellulose membrane (BioRAD) after blocking with BSA and skim milk as described in (Basu and Altuvia 2021) and probed with FLAG M2 monoclonal antibody (Sigma-Aldrich) according to the manufacturer’s protocol. The proteins were visualized using the secondary antibody Anti-Mouse IgG-HRP using Clarity Max™ Western ECL Substrate (BioRAD).

### Northern analysis

RNA samples (10 μg) isolated from strains as indicated were denatured for 10 min at 70°C in 98% formamide loading buffer, separated on 6% acrylamide 8 M urea gels and transferred to Zeta Probe GT membranes (Bio-Rad laboratories) by electroblotting. To detect OxyS, the membrane was hybridized with end labeled OxyS primer (3708) in modified CHURCH buffer (1 mM EDTA, pH 8.0, 0.5M NaHPO4, pH 7.2, and 5% SDS) for 2h at 45°C and washed as previously described (Ben-Zvi et al. 2019). Tm RNA (10Sa) was used as a loading control (1912).

### In vitro RNA synthesis

DNA templates for RNA synthesis: short *mepS* (139 nt) was generated using primers 3720-3721; long *mepS* (248 nt) was generated using 3719-3721; OxyS (109 nt) was generated using primers 2238–2027; OxyS Δ15-33 (94 nt) was generated using primers 2238–2027 The RNAs were synthesized in 50 μl reactions containing T7 RNA polymerase (25 units; New England Biolabs), 40 mM Tris–HCl (pH 7.9), 6 mM MgCl2, 10 mM dithiothreitol (DTT), 20 units RNase inhibitor (CHIMERx), 500 μM of each NTP, and 200 ng of purified PCR templates carrying the sequence of the T7 RNA polymerase promoter. Synthesis was allowed to proceed for 2 h at 37°C followed by 10 min at 70°C. To remove DNA template, 4 U of turbo DNase I (Ambion) was added (37°C, 30 min), followed by phenol/chloroform extraction and ethanol precipitation in the presence of 0.3 M ammonium acetate.

### Primer extension

#### In vivo

Total RNA (30 μg) extracted using Tri reagent (Sigma) from strains as indicated was incubated with end labelled mepS specific primer (3412) at 70 °C for 5 min, followed by 10 min in ice. The reactions were subjected to primer extension at 42°C for 45 min using 1 unit of MMLV-RT (Promega) and 0.5 mM of dNTPs. Extension products were analysed on 6% acrylamide 8 M urea-sequencing gels next to pUC19-MspI labelled marker.

#### In vitro

Annealing mixtures containing in DEPC-treated water, 0.05 pmol of in vitro-synthesized *mepS* RNA, without or with 5 pmol of in vitro-synthesized OxyS RNAs (wild type and mutants) were incubated for 15 min at RT. The mixtures were incubated for another 10 min at RT with 0.6 pmol of end-labeled mepS-specific primer (3740) in 20 mM Tris–HCl, 10 mM magnesium acetate, 0.1 M NH4Cl, 0.5 mM EDTA, 2.5 mM β-mercaptoethanol and 0.5 mM each dNTP upon which reverse transcriptase (Promega; 40 units) was added. cDNA synthesis was allowed to proceed for 7.5 min at 37°C. The extension products were separated on 6% acrylamide 8 M urea-sequencing gels alongside sequencing reactions.

### Survival assays

Overnight cultures of MG1655 *mal*::*lacI*^q^ grown from fresh transformation plates were diluted 1/100 in LB supplemented with the appropriate antibiotics (20 ml in 125ml flasks) and grown at 37°C (200 rpm). IPTG (1mM) to induce OxyS and arabinose to induce *kilR* (0.2%) were added at the time of dilution. Samples to estimate CFU were taken at the indicated time points.

### Mutagenesis assays

To estimate the number of rifampicin resistant mutants ON cultures of wild type and chromosomally encoded *oxyS* mutants were diluted 1/100 in LB and grown shaking at 37°C (200 rpm) to OD600 of 0.1-0.2. Thereafter, cultures treated by 0.2 mM hydrogen peroxide (30 min, 200 rpm) were exposed to 10 mM hydrogen-peroxide for 20 hours (200 rpm). To determine frequencies of mutagenesis, aliquots were taken after 24 h and plated on LB plates containing 100 μg/ml of rifampicin. The numbers of Rif^r^ mutants were normalized to the numbers of viable cells at the 24 h time point.

### Fluorescence microscopy

*E. coli* overnight cultures carrying plasmids were sub-cultured in fresh LB medium supplemented with 1 mM IPTG and Amp. Cultures were grown at 37°C for 3 h before imaging. *Salmonella* cultures carrying *Salmonella* OxyS (P*lac St-oxyS*) were grown for 20 hours in the presence of IPTG. For imaging, cultures were centrifuged and suspended in 20 μl of FM4-64 stain at a final concentration of 100 mg/ml. Samples were adhered using poly-L-lysine coated slides and photographed using Eclipse Ti2 microscope (Nikon, Japan), equipped with Prime BSI camera (Photometrics, Roper Scientific, USA). System control and image processing were performed using NIS-Elements AR Analysis (version 5.30.05; Nikon).

### Phylogenetically weighted sequence logo

The information content in the multiple sequence alignment (MSA) was used to judge the conservation of sRNAs and sRNA promoters. 1340 OxyS homologs including 150 nt upstream of start of sRNA were aligned with MAFFT (Katoh and Standley 2013) (maxiterate 1000, localpair) and the resulting MSA was transformed into a count table storing the appearance of the four nucleotides at each alignment position. Gaps were not counted. To account for the non-uniform organism coverage for different phylogenetic groups the CopraRNA weighting procedure based on 16S rDNA was used. Instead of counting any appearance of a character by 1, it was counted by the weight of the respective organism. The weighted count table was visualized with WebLogo3 (Crooks et al. 2004).

### Phylogenetic tree

A 16S rDNA alignment was done with MAFFT (Katoh and Standley 2013) (retree 2, maxiterate 0), the distance matrix (F81 distance correction) and a maximum likelihood tree was calculated with the phangorn R-package (Schliep 2011). The *Y. pseudotuberculosis* YPIII (NC_010465) 16S rDNA was used as outgroup for rooting.

### Comparative target prediction

For each of the 146 organisms 14 organism from their local phylogenetic neighborhood were selected based on the 16s rDNA-based distance matrix (Figure S2). For each set of 15 organisms a target prediction with CopraRNA was done (Wright et al. 2013; Georg et al. 2020) using the Turner99 energy model.

### Interaction site conservation and clustering

The homologous sRNAs and 5’UTRs were aligned using MAFFT (Katoh and Standley 2013) (maxiterate 1000, localpair). IntaRNA 2.0 (Mann et al. 2017) was used to compute position-wise spot probabilities how likely a combination of positions is covered by an interaction for each sRNA/UTR pair. The resulting probability matrix was mapped to the sRNA and UTR alignments were weighted by the organism specific CopraRNA weight summed up field-wise to get a combined interaction probability landscape. Peak areas, representing potential conserved interaction sites, were identified by the R “contourLines” function. The optimal and the first sub-optimal IntaRNA predicted sRNA/UTR interactions for each organism were assigned to the identified peaks. Interactions assigned to the same peak share the same interaction regions in sRNA and UTR, but they might differ on sequence level or in the actually interacting nucleotide pairs. In a next step all interactions assigned to the same peak are clustered based on their sequence identity and the positions of the interaction pairs. This step relies on a good local alignment of the sub-sequences covered by the peak area. Thus a sub-alignment of the sRNA and UTR sequences is cut from the global alignment based on the peak coordinates and the resulting sequences are re-aligned with DIALIGN-TX (Subramanian et al. 2008).

### Gene set enrichment analysis

Gene ontology annotations were extracted from UniProt (The UniProt Consortium 2023). For organisms not represented in UniProt the *E. coli* annotations were used for homologous genes. The top 200 predicted targets for each organism were subjected to a gene set enrichment analysis with the topGO R-package (Alexa et al. 2006) using the fisher test statistic. All terms with an p-Value ≤ 0.2 were pooled for the conservation analysis. The number of terms with high semantic similarity was reduced with the GOSemSim R-package (https://academic.oup.com/bioinformatics/article/26/7/976/213143). For terms with a sematic similarity of ≥ 0.4 after the “wang”method and an overlap of enriched genes ≥ 50 only the more relevant term was kept. Relevance was scored based on the geometric mean of the information content and the normalized frequency of the term in the investigated organisms.

## ACKNOWLEDGEMENTS

We are grateful to Prof. Sigal Ben-Yehuda for her hep with Fluorescence microscopy. This work was supported by: the Israel Science Foundation founded by The Israel Academy of Sciences and Humanities (138/18), and the Deutsche Forschungsgemeinschaft (grant no. DFG GE 3159/1-1).

## Author Contributions

M.E.-W., F.H., B.S., and J.G. performed the experiments. M.E.-W., F.H., J.G. and S.A. conceived the experiments and analyzed the data. S.A. wrote the manuscript and managed the project.

